# AI-assisted selection of mating pairs through simulation-based optimized progeny allocation strategies in plant breeding

**DOI:** 10.1101/2023.01.25.525616

**Authors:** Kosuke Hamazaki, Hiroyoshi Iwata

## Abstract

Emerging technologies such as genomic selection have been applied to modern plant and animal breeding to increase the speed and efficiency of variety release. However, breeding requires decisions regarding parent selection and mating pairs, which significantly impact the ultimate genetic gain of a breeding scheme. The selection of appropriate parents and mating pairs to increase genetic gain while maintaining genetic diversity is still an urgent need that breeders are facing. This study aimed to determine the best progeny allocation strategies by combining future-oriented simulations and numerical black-box optimization for an improved selection of parents and mating pairs. In this study, we focused on optimizing the allocation of progenies, and the breeding process was regarded as a black-box function whose input is a set of parameters related to the progeny allocation strategies and whose output is the ultimate genetic gain of breeding schemes. The allocation of progenies to each mating pair was parameterized according to a softmax function of multiple selection criteria. Different selection criteria are weighted in the function to balance genetic gains and genetic diversity optimally. The weighting parameters were then optimized by the black-box optimization algorithm called StoSOO via future-oriented breeding simulations. Simulation studies to evaluate the potential of our novel method revealed that the breeding strategy based on optimized weights attained almost 10% higher genetic gain than that with an equal allocation of progenies to all mating pairs within just four generations. Among the optimized strategies, those considering the expected genetic variance of progenies could maintain the genetic diversity throughout the breeding process, leading to a higher ultimate genetic gain than those without considering it. These results suggest that our novel method can significantly improve the speed and efficiency of variety development through optimized decisions regarding the selection of parents and mating pairs. In addition, by changing simulation settings, our future-oriented optimization framework for progeny allocation strategies can be easily implemented into general breeding schemes, contributing to accelerated plant and animal breeding with high efficiency.

## 1 Introduction

To meet the increasing demand for agricultural products caused by the population explosion, developing new cultivars with desired agronomic characteristics, such as high yield, good quality, and efficient nutrient use, is urgently needed (Capper, 2011). However, in plant breeding, for instance, conventional breeding methods require several years or longer to produce new cultivars, even in annual crops, which makes it challenging to meet the increasing food demand in the near future (Eggen, 2012). Genomic selection (GS) is expected to be a critical methodology for accelerating the evaluation and selection of superior genotypes that can be implemented in conventional breeding (Meuwissen et al., 2001; Jannink et al., 2010). In GS, genotypic values of target traits are predicted using genomic prediction (GP) models based on genome-wide marker data, and the predicted values are used for selection in breeding schemes (Meuwissen et al., 2001). GS enables individual-based and crossing-based selection according to the genotypic values predicted by the models with fewer field evaluations of target traits, which leads to highly efficient and rapid breeding (Zhong and Jannink, 2007; Jannink et al., 2010; Crossa et al., 2017; Lehermeier et al., 2017). The superiority of GS over selections based on either phenotypic date or pedigree-derived prediction (pedigree BLUP) has been reported in many simulations and empirical studies (Goddard, 2009; Hayes et al., 2009; Jannink et al., 2010; Goddard et al., 2011; Crossa et al., 2013; Hickey et al., 2014; García-Ruiz et al., 2016; Rabier et al., 2016). However, even though GS has contributed to speeding up breeding during selections, the GS itself does not aim to optimize decisions in breeding. Thus, the ultimate decisions regarding parent selection and choosing mating pairs as the beginning step of the breeding scheme still largely rely on breeders’ experiences because many data-based approaches have not been fully utilized by breeders due to target traits’ complexity, limited resources, or implementation difficulty. In other words, the advent of GS has not yet entirely led to optimized decisions in breeding programs.

To solve the above problems and optimize the decisions regarding (1) selection, (2) mating, and (3) progeny allocation, there has been much discussion on improving the strategies in breeding schemes. First, as for the selection strategy, different selection criteria have been used to increase the efficiency of GS. While various GP models have been developed to improve the accuracy of GS (Meuwissen et al., 2001; Gianola and van Kaam, 2008; VanRaden, 2008; de los Campos et al., 2009; Jannink et al., 2010; Habier et al., 2011; de Los Campos et al., 2013; González-Camacho et al., 2016), genomic estimated breeding value (GEBV), the total sum of estimated additive marker effects, has been generally utilized for GS as a selection criterion (Meuwissen et al., 2001). Although GEBV is usually used for a short breeding period with a few cycles of selection and crossing, it does not guarantee an increased genetic gain in a long-term breeding process (Sonesson et al., 2012; Lin et al., 2017; Akdemir et al., 2019). Because genetic variance is greatly reduced due to the Bulmer effect (Bulmer, 1971; Van Grevenhof et al., 2012) and genetic diversity is also reduced by inbreeding and random drift under truncation selection (Falconer and Mackay, 1996; Li et al., 2008), the genetic gain rapidly reaches a local, rather than global, optimum, i.e., a plateau, in GEBV-based GS cycles (Moeinizade et al., 2019; Zhang and Wang, 2022). Other selection criteria have been proposed to solve the issue of GEBV-based GS by maintaining the genetic diversity and further increasing the genetic gain expected in the long-term breeding program. Weighted genomic estimated breeding value (WGEBV) is a criterion that utilizes marker allele frequencies to emphasize the marker effects of rare alleles (Goddard, 2009; Jannink, 2010). Compared to GEBV, WGEBV can maintain genetic diversity in breeding populations over the long term by suppressing the Bulmer effect (Goddard, 2009; Jannink, 2010). Other selection criteria, such as optimal haploid value (Daetwyler et al., 2015), with an expected maximum haploid breeding value (Müller et al., 2018), and optimal population value (Goiffon et al., 2017), have been introduced to maintain genetic diversity by including haplotype information and have indeed improved the final genetic gains at the end of the breeding scheme in long-term breeding programs compared to those from GEBV-based GS. Although these selection criteria are helpful in selecting parental candidates, they do not directly enable the automatic optimization of decisions regarding the selection of parents. Therefore, breeders must make ultimate decisions based on these criteria by themselves. Also, an optimal contributions selection (OCS) method, which aims to optimize decisions regarding parent selection, was proposed to help breeders decide on more appropriate breeding strategies rather than just use selection criteria (Meuwissen, 1997; Grundy et al., 1998). OCS aims to optimize the expected contribution vector, i.e., the number of times each individual is used as a parent for the next generation, by maximizing the genetic gains while constraining the inbreeding in the next generation. Although the efficiency of the OCS-based GS was evaluated in several simulation studies (Gorjanc et al., 2018; Cowling et al., 2023), OCS still cannot assist in identifying optimal mating pairs even for the next generation.

Second, as for the problems regarding the selection of appropriate mating pairs, i.e., the mating design problem, many studies in animal breeding have long been conducted to select optimal pairs while considering mating constraints (Jansen and Wilton, 1985; Kinghorn, 2011), but they mostly lacked a long-term perspective. Then, a look-ahead selection (LAS) approach has been proposed as a method for simultaneously optimizing decisions regarding the selection of appropriate parents and mating pairs in plant breeding (Moeinizade et al., 2019). LAS can optimize whether each pair of individuals should be mated to maximize the final genetic gain via look-ahead simulations under two simplified assumptions: one progeny is obtained from each mating pair, and random mating occurs after the first selection and mating cycle. Although this approach achieved higher genetic gains than the previous selection criteria in the final generation by considering optimal mating pairs, the assumptions were too simple to estimate the final genetic gain in practical breeding programs, and LAS could not optimize the number of progenies allocated to each mating pair, i.e., progeny allocation strategy.

Thirdly, as for the progeny allocation strategy, only one study has attempted to optimize the allocation strategy (Hunter and McClosky, 2016) although it is known to have a significant impact on the ultimate genetic gain (Wang et al., 2018). In their study, the number of progenies allocated to each pair was optimized by searching for the Pareto surface in the plane formed by the mean and variance of progenies in the next generation (Hunter and McClosky, 2016), utilizing the multi-objective optimal computing budget allocation method (Lee et al., 2010). However, this method lacks the future-oriented perspectives required for the mid-term/long-term breeding process. Moreover, there is no study on the allocation optimization of progenies for the final genetic gain under multiple selection and crossing cycles, although a couple of previous studies tried to optimize breeding program decisions from other viewpoints (Amini et al., 2021; Diot and Iwata, 2022; Moeinizade et al., 2022).

As described so far, regardless of the duration of breeding schemes, i.e., short-term or long-term, how decisions are made significantly impacts the ultimate outcome of the schemes when parent selection and mating are repeated multiple times. Thus, if the optimization framework regarding the selection of parents and mating pairs is achieved, it can be widely applied to various types of breeding programs, i.e., small-scale and large-scale breeding programs. As for small-scale breeding programs with limited resources, particularly in plant breeding, since many minor crops, including neglected and underutilized species (NUS), have attracted much attention as breeding targets, there is increasing demand for realizing more efficient breeding in small-scale programs for minor crops (Padulosi et al., 2013; Kamenya et al., 2021). Even if NUS has attracted little attention and has been entirely ignored by plant breeders so far, it is now expected to have the potential to diversify agriculture and address climate change. Another example of small-scale breeding is breeding schemes promoted by small and medium enterprises (Zambon et al., 2019). Since there are many agricultural small and medium enterprises in Asian countries, optimizing decision-making in small-scale breeding will lead to increased profits in the agricultural market. From these viewpoints, our study, which helps breeders’ decisions on hybridization, is essential to improve breeding efficiency not only in large-scale but also small-scale breeding programs.

The overall objective of the study was to go beyond the conventional selection criterion-based GS by optimizing a progeny allocation strategy to maximize the ultimate genetic gains after multiple cycles of selection of parents and mating pairs. There are two significant issues in this optimization. The first issue is that it is challenging to describe the breeding process explicitly because the future state of a breeding population is unknown. The second issue is that breeding is a stochastic rather than deterministic process since progenies are randomly produced in each generation through meiosis. Therefore, the specific objectives of this study were (1) to evaluate the final genetic gain via future-oriented breeding simulations and (2) to achieve an optimal mating pair selection via a numerical optimization approach conducted by an artificial intelligence (AI) breeder while addressing the two issues above. First, GS combined with future-oriented breeding simulations enables us to evaluate the final genetic gain in a breeding scheme with multiple cycles, leading to a solution to the first issue. Second, the AI breeder in this study referred to a virtual breeder in our computer who can optimize the decisions regarding the progeny allocation strategy by using both the breeding simulations and some optimization algorithm. In this study, to design this AI breeder, we applied black-box optimization by considering the breeding process achieved by the above simulations as a black-box function whose input is a set of parameters for the allocation strategy and whose output is the ultimate genetic gain. We employed a black-box optimization algorithm, StoSOO (Valko et al., 2013), which can also optimize a stochastic black-box function, such as a breeding process, presenting a solution to the second issue. Multiple selection criteria are used to allocate progenies, and the optimized progeny allocation can automatically determine the optimal combination of criteria and weights to be used in the allocation. Further, the optimized progeny allocation can help breeders select the appropriate mating pairs because the AI breeder can offer information on which cross combination is prospective by showing the optimized number of progenies in continuous values. In this study, we mainly focus on optimizing the progeny allocation strategy as a proof of concept. Still, by changing the simulation settings and the parameters to be optimized, we can easily extend our framework to assist breeders in making decisions in different types of hybridization, such as parental selection in bi-parental/multi-parental populations.

## 2 Materials and methods

### 2.1 System design

We assumed that breeders proceed with a breeding scheme according to the progeny allocation strategy proposed by an “AI breeder”, a novel decision-making system in this study (Figure 1A). The breeders can obtain a set of parameters representing the optimal allocation strategy,{**ĥ**^(*τ*)^}*τ*=0,1,2,3 passing the information on a parent panel, i.e., marker genotype, genetic marker effects, and recombination rates between genetic markers, to the AI breeder. Here, we briefly describe the overview of the framework in which the AI breeder optimizes the strategy for the resource allocation of progenies.

**Figure 1.**
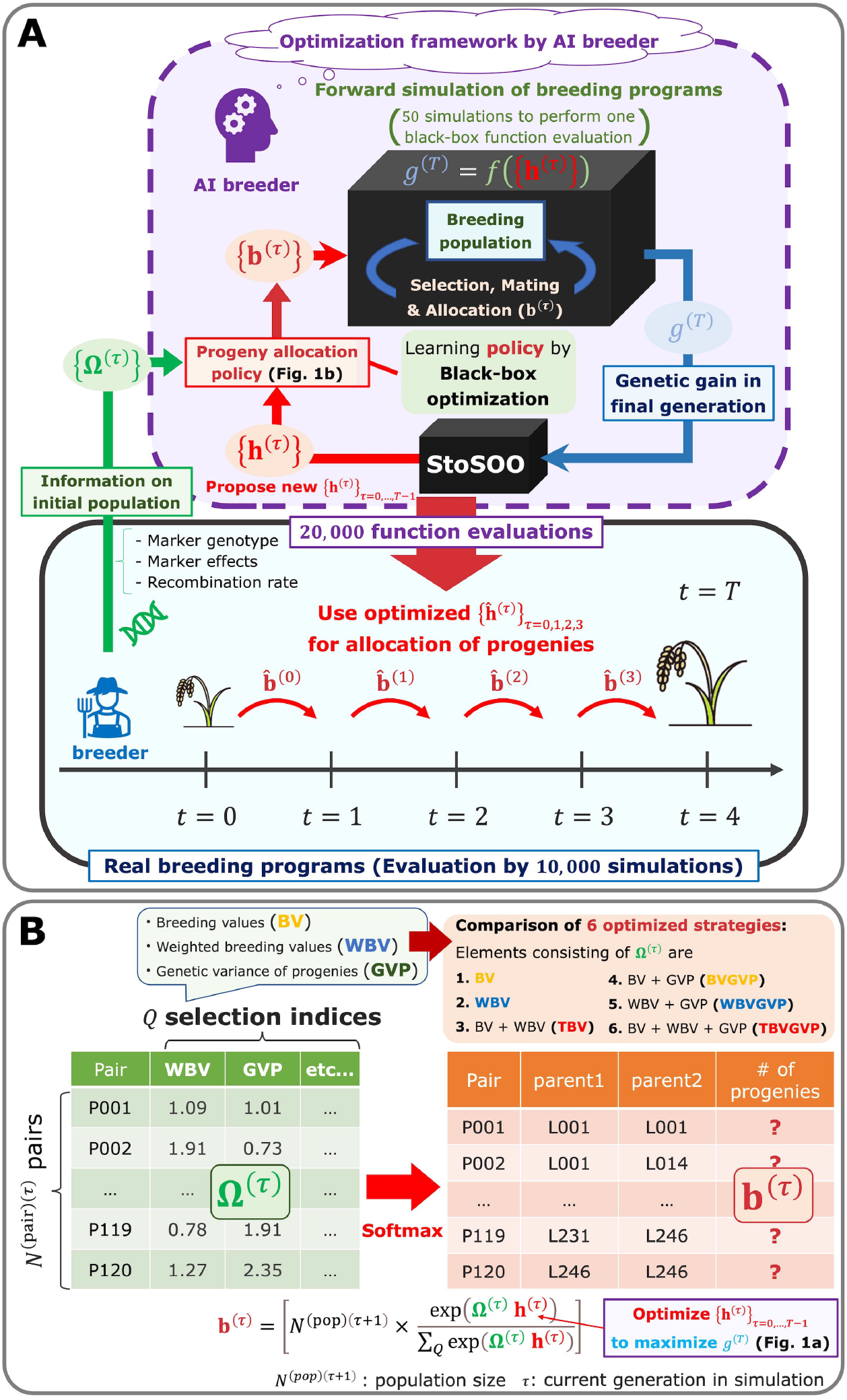
System design assumed in this study. (**A**) The framework to optimize decision-making in the breeding scheme. (**B**) The resource allocation strategy of progenies developed in this study.

After receiving the parent panel information, the AI breeder attempts to simulate breeding schemes by defining the allocation strategy (Figure 1A). At each generation *τ* in the simulated schemes (*τ*∈ {0,1,2,3} is a current generation number in simulations conducted by the AI breeder, whereas *t* ∈{0,1,2,3} appeared in the subsequent subsections is a current generation number in breeding schemes proceeded by actual breeders), the breeder first computes the weighted breeding values (WBV) to select the parent candidates from the current generation. Then, after determining mating pairs by the diallel crossing between the parent candidates, given the parameter **h**^(*τ*)^, the breeder automatically determines the number of progenies allocated to each pair **b**^(*τ*)^, by utilizing the information on selection criteria Ω ^(*τ*)^ computed from the marker genotype and effects, such as breeding values (BV), WBV, and expected genetic variance of progenies (GVP) (Figure 1B). In other words, we assumed that **b**^(*τ*)^ is determined by applying the softmax function to the weighted sum of the selection criteria, as shown in Equation 1.

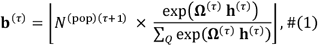

where *N*^(pop)(*τ*+1)^ is the population size in the next generation *τ*+1, and ⌊*x* ⌋is the floor function of *x* (i.e., the maximum integer that is equal to or lower than *x*). The softmax function is used to allocate progenies in proportion to probabilities reflecting the weighted sum of the selection criteria. Here, **h**^(*τ*)^ can be interpreted as the weighting parameters that correspond to the importance of each selection criterion. After determining **b**^(*τ*)^, the AI breeder can simulate the next generation based on the principle of meiosis. Then, by repeating the parent selection and mating steps toward the final generation of the breeding scheme, the AI breeder evaluates the performance of the allocation strategy with **h**^(*τ*)^. Thus, the goal of the AI breeder is to estimate {**h**^(*τ*)^} _*τ* =0,1,2,3_ that maximizes the final genetic gain via simulations of the breeding process. To optimize the weighting parameters, a black-box optimization algorithm called StoSOO (Valko et al., 2013) was used in this study (Figure 1A). The StoSOO algorithm is an extension of the SOO algorithm (Munos, 2011; Preux et al., 2014), which performs global optimization of deterministic black-box functions based on a tree-based search and can be applied to stochastic black-box functions. Using the StoSOO algorithm with a finite number of function evaluations can provide a global quasi-optimal solution of an implicit stochastic function within the number of evaluations. To apply StoSOO to our problem, the AI breeder performs 50 breeding simulations with one set of {**h**^(*τ*)^} _*τ* =0,1,2,3_, and the empirical mean of the 50 final genetic gains is regarded as the objective function for StoSOO. The function evaluations are repeated 20,000 times in StoSOO, and a set of optimized parameters {**ĥ**^(*τ*)^} _*τ* =0,1,2,3_ is passed from the AI breeder to the real breeders.

In this study, we evaluated this novel optimization framework by conducting simulation studies as a proof of concept. We prepared six versions of the optimized allocation strategy according to the components of **Ω**^(*τ*)^ as follows: (a) only BV, (b) WBV, (c) both selection criteria (i.e., BV and WBV), (d) expected GVP in addition to BV, (e) expected GVP in addition to WBV, and (f) expected GVP in addition to (c) (Figure 1B). We compared these six optimized strategies to (g) the equal allocation strategy that equally allocates progenies to each pair, that is, **h**^(*τ*)^ = **0**,by conducting 10,000 breeding simulations using the optimized {**ĥ**^(*τ*)^} _*τ* =0,1,2,3_ (Figure 1A). In the following results, we abbreviate the above strategies as (a) BV, (b) WBV, (c) TBV (Two BVs), (d) BVGVP, (e) WBVGVP, (f) TBVGVP, and (g) EQ. Here, for the BV and WBV strategies, we only used one criterion for allocating progenies. For simplicity, we also assumed true marker effects to compute the selection criteria. In other words, true QTL positions and effects were known throughout the study. The problem of using the true marker effects will be discussed in the **Discussion** section.

### 2.2 Overview of the methods

This section provides an overview of the method used in this simulation study (Figure S1). We first simulated genome-wide markers and QTLs for the parent panel and the corresponding QTL effects. The selection of parent candidates, determination of mating pairs, and allocation of progenies to each mating pair were then repeated in sequence to move forward with a breeding scheme until the final generation. Selection and mating processes were carried out based on two selection criteria: BV and WBV, and the expected GVP. We compared two breeding strategies regarding the allocation of progenies: equal allocation and allocation optimized by a black-box optimization algorithm, StoSOO. These strategies were evaluated by simulating 10,000 breeding schemes based on each strategy.

Each element of the simulation study is described in the following subsections. First, the two selection criteria, BV and WBV, are described, followed by the simulation of genome-wide markers and QTLs, the selection and mating processes in breeding schemes, and the evaluation of breeding schemes. Because optimized allocation requires the simulation and evaluation of breeding schemes, the method of evaluating breeding schemes is explained before describing the optimized allocation strategy. The calculation of the GVP required for optimized allocation is presented at the end of this section.

### 2.3 Selection criteria

We used two selection criteria, BV (Meuwissen et al., 2001) and WBV (Goddard, 2009; Jannink, 2010), to select parent candidates and allocate progenies in breeding schemes since these are the two typical breeding values (Figure S1 (1)). Here, we only used WBV for the selection step to maintain the diversity, but we used both criteria for the allocation step.

The first criterion, BV, can be expressed as in Equation 2:

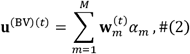

where *M* is the number of markers, *N*^(pop)(*t*)^ is the number of genotypes for a population with generation *t* in a breeding scheme, **u** ^(BV)(*t*)^ is an *N*^(pop)(*t*)^ × **1** vector of BV for generation *t* 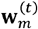 is an *N*^(pop)(*t*)^ × **1** vector of marker genotype at *m*-th marker with scores of -1,0, or 1for generation *t* and *α*_*m*_ is a true marker effect at the *m*-th marker. In this study, we assumed that the true marker effect *α*_*m*_ was known. BV is a criterion for expressing additive genotypic values directly transmitted to the next generation and is often used in conventional breeding schemes with GS (Meuwissen et al., 2001).

The second selection criterion, WBV, can be expressed as in Equation 3:

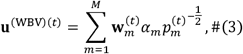

where **u**^(WBV)(*t*)^ is an *N*^(pop)(*t*)^ × **1** vector of WBV for generation *t*, 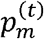 is the allele frequency at the *m*-th marker for generation *t*, and the other terms are defined in Equation 2. Emphasizing the effects of rare alleles, as in Equation 3, leads to the successful maintenance of genetic diversity, thus WBV can benefit the genetic gain in long-term breeding programs compared to BV (Goddard, 2009; Jannink, 2010).

### 2.4 Simulation of the parent panel and genome structure

Genome-wide markers and QTLs for founder haplotypes were generated using the coalescent simulator GENOME (Liang et al., 2007) (Figure S1 (2)). We assumed a diploid with the number of chromosomes ten (2*n*=20), and that there were 500 loci, including both markers and QTLs, on each chromosome. At first, 4,000 founder haplotypes for each chromosome were independently generated using GENOME with the following parameters: “-pop 1 4000 -N 4000 -c 1 -pieces 100000 -rec 0.0001 -s 4000 -maf 0.01 -tree 0 -mut 0.00000001”. Among the 4,000 loci, 500 loci whose minor allele frequency (MAF) was equal to or larger than 0.01 were randomly selected. Subsequently, 2,000 founders were generated by sampling two haplotypes from all founder haplotypes with replacement. These founders were randomly mated for one generation to simulate the pre-breeding materials of the 2,000 genotypes. These procedures for simulating the genetic resources are similar to those in the R package “BreedingSchemeLanguage” by Yabe et al. (Yabe et al., 2017). From these simulated genetic resources, *N*^(pop)(0)^= 250 g enotypes were selected for the parent panel of a breeding scheme so that these 250 genotypes represented the simulated genetic resources in terms of genetic diversity. We applied the k-medoids method to the marker genotypes of the pre-breeding material using the “pam” function of the R package “cluster” version 2.1.2 (Maechler et al., 2021). In the k-medoids method, after clustering the pre-breeding materials into 250 groups, we selected the medoids as the representative genotypes from each group. This parent panel was regarded as generation *t*=0 in a breeding scheme. Here, the parent panel consisted of a relatively small number of genotypes and was also genetically close to founders compared to usual breeding populations used in programs for major crops. These simulation settings, including the assumption for the parent panel, are a little apart from situations in large-scale breeding for major crops but can be justified when we assume the breeding schemes for the NUS whose breeding has not been promoted in the past.

### 2.5 Simulation of QTLs and genotypic values

For each chromosome, 400 loci were randomly sampled from the 500 loci and were regarded as single nucleotide polymorphism (SNP) markers available for breeders. In simulating QTLs and phenotypes, we assumed quantitative traits with a simple genetic architecture as target traits, as described below (Figure S1 (3)). First, among the remaining 100 loci, 2 loci per chromosome were randomly selected as QTLs (i.e., a total of *M*^(QTL)^ =20 QTLs). This number of QTLs was determined assuming the situation where quantitative traits controlled by the relatively small number of QTLs still remain as target traits in small-scale breeding for the NUS. QTL effects were then sampled from the normal distribution, as shown in Equation 4.

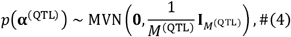

where **α**^(QTL)^ is an *M*^(QTL)^ ×1 vector of QTL effects and 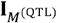 is an *M*^(QTL)^ ×*M*^(QTL)^ identity matrix. Here, QTL effects were assumed to be additive for simplicity. The true additive genotypic values **u**^(TGV)(0)^ for the parent panel were then simulated using Equation 2, where each marker was replaced by each 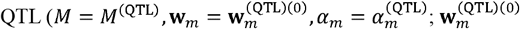, is an *N*^(pop)(0)^ × **1** vector of genotype scores at *m*-th QTL for the parent panel). Thus, **u**^(TGV)(0)^ = **u**^(BV)(0)^ when the true marker effect α_*m*_ is known as the setting in this study. Since we used the true marker effects, a heritability was assumed to be 1 throughout the study.

### 2.6 AI-assisted breeding scheme

The breeding scheme in this study was carried out according to the following steps (Figure S1 (4)). Here, we assumed small-scale plant breeding programs with recurrent selection for simplicity and due to the computational limitations, but changing the settings of the following breeding scheme will lead to the extension of our framework for various types of breeding programs.

#### 1 Select parent candidates (Figure S1 (4-1))

Based on the order of increasing WBV computed using the true marker (QTL) effects **α**^(QTL)^ for the current generation *t* (*t* ∈{0,1,2,3}, the top *N*^(sel)(*t*)^ genotypes were selected as parent candidates for the next generation. In this study, we assumed *N*^(sel)(*t*)^ was constant over generations, i.e., *N*^(sel)(*t*)^= *N*^(sel)^ =15. We used only WBV for the selection step in this study because selecting parents based on BV often failed to maintain the genetic diversity of the population when we chose a high selection intensity in the optimized allocation strategy.

#### 2 Determine mating pairs for the next generation (Figure S1 (4-2))

From the selected *N*^(sel)(*t*)^ parent candidates, diallel crossing, including selfing, was assumed to determine the mating pairs for the next generation, that is, 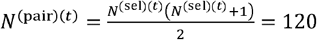 pairs were prepared for crossing.

#### 3 Allocate progenies to each mating pair (Figure S1 (4-3))

The following two strategies were used to allocate progenies to each mating pair determined in Step 2, and then a crossing table was created. Both strategies generated *N*^(pop)(*t*+1)^ progenies for the next generation. In this study, we assumed *N*^(pop)(*t*+1)^ was also constant over generations, i.e., *N*^(pop)(*t*+1)^ =*N*^(pop)^ =250.

##### a The optimized allocation method

One strategy was the optimized allocation method developed in this study, which applied the softmax function to the weighted sum of the selection criteria computed in Step 1. For more details, please refer to the **Progenies allocation strategies** subsection.

##### b Equal allocation method

We also prepared an equal allocation method that allocated *N*^(pop)(*t*+1)^ progenies equally to the *N*^(pair)(*t*)^ mating pairs. Thus, three progenies were allocated to each of the 10 mating pairs, and two progenies were allocated to each of the remaining 110 mating pairs.

#### 4 Generate progenies for the next generation (Figure S1 (4-4))

According to the crossing table created in Step 3, two gametes were generated for each mating pair, considering the recombination between markers. Recombination rates between markers were computed using the Kosambi map function (Kosambi, 1943; Zhao and Speed, 1996) based on the linkage map generated by GENOME. These two gametes were then combined to create one new progeny, resulting in a new population (with generation *t*+1) of *N*^(pop)(*t*+1)^ progenies.

#### 5 Repeat the parent selection and mating process

Steps 1-4 were repeated four times until the population reached the final generation, i.e., *t* =*T* − 1where *T* is the final generation number.

Finally, the results for each strategy were evaluated using the true additive genotypic values **u**^(TGV)(T)^ for the latest population with generation *T*. Here, the final generation was set as *T*=4 in this study, which was determined assuming the situation where we needed to quickly achieve high genetic gains in small-scale breeding schemes as one example.

### 2.7 Evaluation of the outcome of breeding schemes

In one breeding scheme, the true additive genotypic value **u**^(TGV)(*t*)^ was computed for each generation *t* (Figure S1 (5)). Then, the top *N*^(top)(*t*)^=5 genotypes were chosen in the order of increasing **u**^(TGV)(*t*)^, and the empirical mean of the **u**^(TGV)(*t*)^ for these *N*^(top)(*t*)^ genotypes, *u*^(top)(*t*)^, was computed to evaluate the population maximum for the breeding strategy. Here, we chose *N* ^(top)(*t*)^ =5 genotypes for the evaluation to represent the top individuals for developing a new variety and to control the stochastic variation between simulations to some extent. For a given simulation dataset of the parent panel, we conducted 10,000 different breeding schemes for each strategy and evaluated the empirical mean of the *u*^(top)(*t*)^ using these 10,000 simulation results. The simulation and its evaluation were repeated for 10 replicates of the phenotype simulation with different QTL positions and effects.

### 2.8 Progenies allocation strategies

In this study, we developed a novel breeding strategy to quasi-optimize each selection criterion regarding the allocation of progenies (Figure S1 (6) and Figure 1b). In other words, we determined the number of progenies allocated to each mating pair at generation *τ*,**b** ^(*τ*)^, by applying the softmax function to the weighted sum of the selection criteria instead of the equal allocation strategy described above, as in Equation 1.

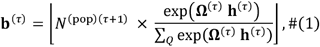

where **b**^(*τ*)^ is an *N*^(pair)(*τ*)^×1 vector representing the number of progenies allocated to each mating pair at generation ×, Ω ^(*τ*)^ is an *N*^(pair)(*r*)^× matrix×*Q* of *Q* different selection criteria for an average progeny of each mating pair (e.g., BV, WBV, or the genetic variance of progenies in a subsequent generation (GVP) for each pair), and ⌊ *x* ⌋ is the floor function (i.e., the maximum integer that is equal to or lower than *x*). In the softmax function, we can convert a vector of some weighted goodness for each mating pair, Ω^(*τ*)^ **h**^(*τ*)^, to a vector of probabilities reflecting the relative value of each element by allocating more progenies to better pairs. Here, **h**^(r)^ is a *Q* ×1 vector required for determining **b**^(*τ*)^, which corresponds to how much importance is given to each selection criterion. Since the final genetic gain, *g*^(*T*)^, can be represented as the output of the black-box function *f* whose input is a set of **h**^(*τ*)^, i.e., *g*^(*T*)^ =*f* ({**h**^(*τ*)^} _*τ* =0,1,2,3_), the main goal of this strategy is to estimate **h**^(*τ*)^ that maximizes *g*^(*T*)^ via breeding simulations and the black-box optimization algorithm. When **h** ^(*τ*)^ =0, the softmax allocation strategy based on Equation 1 is the same as the equal allocation strategy. The final genetic gain was defined as the improvement in the true genotypic values of the top *N*^(top)(*t*)^ = 5 genotypes of the final population compared with those of the parent panel, that is, *g*^(*T*)^ =*u* ^(top)(*T*)^ -*u*^(top)(0)^.

### 2.9 Optimization of h^(*τ*)^ to maximize the final genetic gain

In this study, we used the StoSOO algorithm (Valko et al., 2013) to optimize the parameter vector **h**^(*τ*)^ (Figure S1 (7)). The StoSOO algorithm is an extension of the SOO algorithm (Munos, 2011; Preux et al., 2014), which performs global optimization of deterministic black-box functions based on a tree-based search and can be applied to stochastic black-box functions. The StoSOO algorithm with a finite number of function evaluations can provide a global quasi-optimal solution of an implicit stochastic function within the number of evaluations. To apply StoSOO in our study, we performed 50 simulations using one set of {h^(*τ*)^} ^*τ* =0,1,2,3^, and the empirical mean of the final genetic gains *g*^(*T*)^for these 50 simulations was used as the objective function for StoSOO. Here, we defined the domain of definition as and set as an initial parameter for each element of {h^(*τ*)^} ^*τ* =0,1,2,3^. The function evaluations were repeated 20,000 times, giving us quasi-optimal solutions,{ **ĥ**^(*τ*)^}^*τ*^, for this optimization problem. We then performed 10,000 simulations based on the estimated { **ĥ**^(*τ*)^}^*τ*=0,1,2,3^ to evaluate each optimized strategy developed in this study, as described in the **Evaluation of results** subsection.

### 2.10 Selection criteria and genetic diversity of progenies as the candidates for Ω ^(*τ*)^

As the candidates for **Ω** ^(*τ*)^ to compute b^(*τ*)^, we used the two selection criteria, BV and WBV, and the expected GVP for each mating pair (Figure S1 (8)). The BVs and WBVs of the parent candidates were computed using Equations 2 and 3, and the mean of the selection criteria of the two parents for each mating pair was used as one column vector of **Ω** ^(*τ*)^. This vector, representing BV or WBV for each mating pair, was the same as that for the average progeny of each mating pair in this study because we only assumed additive QTL effects. To compute the GVP, we used Equation 5, based on the idea proposed by Lynch and Walsh (Lynch and Walsh, 1998; Zhong and Jannink, 2007; Lehermeier et al., 2017):

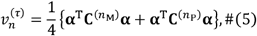

where 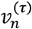 is an expected GVP for mating pair *n* with generation *τ*, **α** is a vector of the true marker effects, 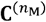 and 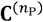 are *M* × *M* covariance matrices caused by the segregation of maternal parent *n*_M_ and paternal parent *n*_P_ of mating pair *n*, respectively. A diagonal (*m, m*) element of 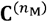, 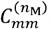, is 0 if an *m*-th marker of *n*_M_ is homozygote, and 1 if otherwise (i.e., heterozygote), and a nondiagonal (*m*_1_, *m*_2_) element of 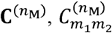, is 0 if the *m*_1_-th or *m*_2_-th marker of *n*_M_ is homozygote, 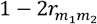 if both *m*_1_-th and *m*_2_-th markers are heterozygote without recombination, and 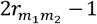 if otherwise (i.e., heterozygote with recombination). Here, 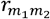 is the recombination rate between The *m*_1_-th and *m*_2_-th markers. In this study, after computing the expected GVP for all mating pairs,**v** ^(*τ*)^ was used as one column vector of **Ω** ^(*τ*)^.

We prepared six different strategies for the components of **Ω** ^(*τ*)^and compared them to the non-optimized strategy as written in the **System Design** subsection, i.e., (a) BV, (b) WBV, (c) TBV, (d) BVGVP, (e) WBVGVP, (f) TBVGVP, and (g) EQ. Even when we used only one criterion for the allocation of progenies in BV and WBV strategies, we optimized the weighting parameter {**h**^(*τ*)^}_*τ* = 0,1,2,3_ by focusing on the optimization of the gradient of the number of progenies allocated to each pair against the relative value of that criterion.

## 3 Results

### 3.1 Evaluation of convergence conditions of the StoSOO algorithm

For the optimization of the allocation strategy to work, it was necessary to confirm that the solution obtained from the algorithm used for optimization, the StoSOO algorithm, converged stably. Thus, we first evaluated the convergence of the solution using the StoSOO algorithm for each strategy. As described above, the AI breeder conducted 50 simulations of the breeding scheme for one function evaluation and repeated 20,000 function evaluations to obtain parameter sets that were quasi-optimized by the StoSOO algorithm. We recorded the quasi-optimized parameter sets for each evaluation step of the objective function and evaluated the corresponding function values. For each of the six optimized allocation strategies, the change in the function values was plotted and compared with the function value based on the equal allocation strategy (Figure S2).

For all strategies on the components of **Ω** ^(*τ*)^, the function value increased almost monotonically and reached the quasi-optimized function value after the 20,000 evaluations, which was much larger than that under the equal allocation strategy (Figure S2). These results indicated that StoSOO successfully improved the expected final genetic gain evaluated by the simulations and optimized the parameter set {**h**^(*τ*)^}_*τ* = 0,1,2,3_, resulting in the superiority of the optimized allocation strategy over the equal allocation strategy.

### 3.2 Genetic gains over four generations

For all strategies on the components of **Ω** ^(*τ*)^, we compared the optimized and equal allocation strategies by plotting the change in the genetic gain for each generation (Figure 2A). We then defined genetic gain in each generation as the difference between the empirical means of the true genotypic values of the top *N*^(*top*) (*t*)^ = 5 genotypes in the current generation (generation *t*) and the parent panel (generation 0). Using the optimized allocation strategies obtained after 20,000 function evaluations, we evaluated the expected genetic gains of the seven strategies, including the six optimized and the equal allocation strategies, based on 10,000 breeding simulation results using one replication for the quantitative phenotype simulation.

**Figure 2.**
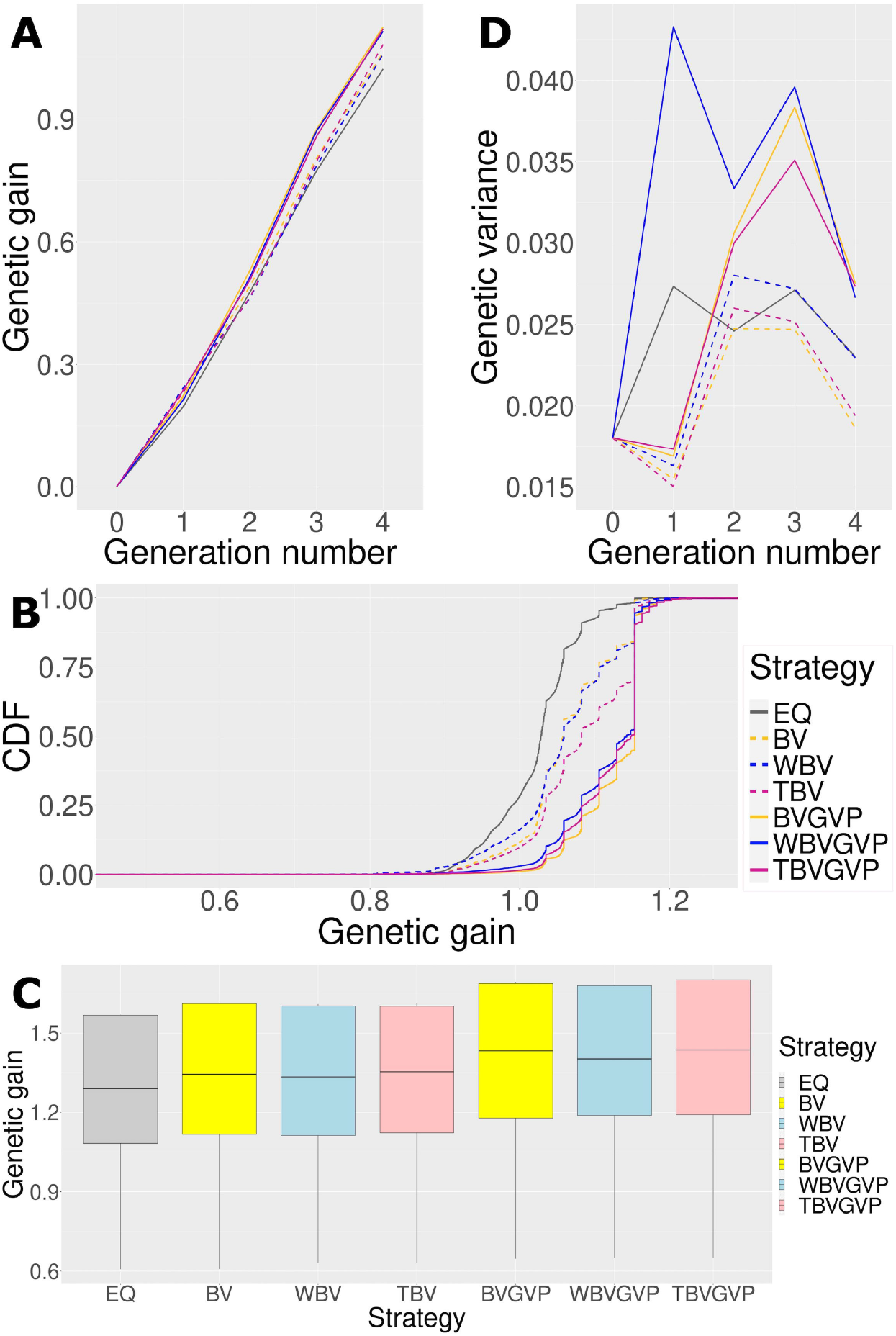
Comparison of the seven strategies (six optimized and one non-optimized) in four ways. (**A**) Change in the expected genetic gains over four generations. (**B**) Genetic gains across different simulation repetitions. (**C**) Genetic gains in the final generation using ten replications for the phenotype simulation. (**D**) Change in the genetic variances over four generations. In all Figures, we compared the seven strategies: EQ: equal allocation (ABD: black solid, C: grey), BV: optimized allocation based on BV (ABD: orange dashed, C: yellow), WBV: optimized allocation based on WBV (ABD: blue dashed, C: light blue), TBV: optimized allocation based on BV and WBV (ABD: red dashed, C: light pink), BVGVP: optimized allocation based on BV and GVP (ABD: orange solid, C: yellow), WBVGVP: optimized allocation based on WBV and GVP (ABD: blue solid, C: light blue), and TBVGVP: optimized allocation based on BV, WBV, and GVP (ABD: red solid, C: light pink). In Figures A and D, generations 0 and 4 correspond to the populations in the initial and final generations, respectively. In addition, generations 1–3 correspond to populations in the middle of the breeding scheme. In Figure B, the horizontal axis shows the final genetic gain of an individual, whereas the vertical axis represents the percentile of the simulation repetitions. As shown in Figure C, for each optimized strategy, the weighting parameter was optimized based on 4,000 function evaluations.

In the final generation, *t* = 4, all the strategies with optimized allocations showed higher genetic gains than the equal allocation strategy (Figure 2A). Strategies that considered GVP outperformed those without GVP in the final generation, while the strategies that only considered the selection criteria BV and WBV showed little difference in genetic gains.

In the first generation, *t* = 1, the strategies that considered GVP showed lower genetic gains than those without GVP (Figure 2A). After the second generation, *t =* 2, strategies with GVP overtook those without GVP, increasing their advantage over the other strategies through the final generation.

### 3.3 Genetic gains across different simulation repetitions

To assess the characteristics of each strategy in more detail, we computed the cumulative distribution functions (CDFs) of genetic gain in the final generation, *t* = 4, for each strategy based on 10,000 simulation repetitions. In Figure 2B, the horizontal axis represents the final genetic gain of an individual, whereas the vertical axis represents the percentile of simulation repetitions. Because the 1^st^ and 99^th^ percentiles correspond to the worst and best performances within the 10,000 simulation repetitions, respectively, the more a CDF curve of a strategy moves toward the right and downward in the graph, the better the performance of the strategy is.

Again, the optimized strategies outperformed the equal allocation strategy; among the optimized strategies, those with GVP showed better performance than those without GVP in terms of the CDFs (Figure 2B). The CDF curves for the optimized strategies were parallel to the vertical axis at the location where the genetic gain was approximately 1.15, suggesting a local optimum within the simulation repetitions. In the optimized strategies with GVP, about 50 % of the breeding simulations achieved genetic gains above this local optimum (BVGVP, 55.1 %, WBVGVP, 47.6 %, TBVGVP, 49.4 %), which were much higher than those in the equal allocation strategy (EQ, 1.85 %) and strategies without GVP (BV, 16.0 %, WBV, 16.4 %, TBV, 30.1 %).

### 3.4 Genetic gains in the final generation using ten replications of the phenotype simulation

We also evaluated the genetic gains in the final generation, *t* = 4, using ten replications of the phenotype simulation to confirm that the optimized strategies showed better results than the equal allocation strategy for the traits with different quantitative trait loci (QTL) positions and effects (Figure 2C). When simulating phenotypes for different traits, we changed QTL positions and their effects, but the number of QTLs and distribution of QTL effects were fixed. Also, the heritabilities were assumed to be 1 for all ten replications since the true marker effects were utilized for the optimization.

To reduce the computational time for the optimization in evaluating ten replications of the phenotype simulation, optimized (or quasi-optimized) allocation strategies were determined after 4,000 function evaluations. The number of function evaluations was determined to ensure that the function value reached a certain level, although it did not converge, and to reduce the computation time based on the results in the subsection that evaluated the convergence conditions of the StoSOO. Here, the StoSOO algorithm does not necessarily have to be converged since the evaluation for the ten replicates of phenotype simulation was conducted to prove that the optimization algorithm could improve the genetic gain to some extent compared to the non-optimized strategy.

Even when the QTL positions and effects of the target trait were changed, the optimized strategies consistently outperformed the equal allocation strategy, and the strategies with GVP outperformed those without GVP (Figure 2C). When we evaluated the improvement rate in the final genetic gain of each optimized strategy compared to the equal allocation strategy using ten replications of the phenotype simulation, a similar trend was observed between the strategies with and without GVP (Table 1). In addition, the optimized strategies at least improved the genetic gain compared to the equal allocation for all ten replications. In particular, the final genetic gains for TBVGVP improved by 7.22–12.4 % compared to those for the equal allocation strategy within just four generations (Table 1).

**Table 1.** Minimum, mean, and maximum improvement rate in the final genetic gain of the optimized strategies compared to equal allocation strategy among ten different target traits (%)

| Method | Min (%) | Mean (%) | Max (%) |
| --- | --- | --- | --- |
| BV | 0.00 | 2.97 | 4.89 |
| WBV | 2.07 | 2.88 | 3.89 |
| TBV | 2.11 | 3.52 | 5.86 |
| BVGVP | 6.40 | 8.63 | 12.4 |
| WBVGVP | 6.99 | 8.28 | 9.95 |
| TBVGVP | 7.22 | 9.33 | 12.4 |

### 3.5 Genetic diversity over four generations

We compared the optimized and equal allocation strategies in genetic diversity across generations (Figure 2D). The genetic diversity of a population in each generation was calculated as the genetic variance of the true genotypic values in the population.

First, in *t* =1, all the optimized strategies, except WBVGVP, showed lower genetic variances than those in the parent panel, *t* =0 (Figure 2D). From *t* =2 to *t* =4, genetic diversity was preserved in all strategies compared to *t* =0. In these generations, strategies with GVP maintained higher genetic variance than those without GVP and the equal allocation strategy. The difference between the strategies with or without GVP was most observed at *t* =3, followed by a sharp decrease in genetic variance for the former strategies through the final generation, *t* =4.

In contrast, strategies using different selection criteria (BV or WBV) showed little difference in genetic variance (Figure 2D). The strategy using only WBV as a selection criterion appeared to maintain a higher genetic variance than those using BV or both BV and WBV. However, the difference was much smaller than between strategies with or without GVP.

### 3.6 Optimized weighting parameters for each strategy

After the 20,000 function evaluations, we obtained a set of optimized weighting parameters {**h**^(*τ*)^}_*τ* = 0,1,2,3_for each allocation strategy (Tables 2 and Tables S1 and S2). A large value of **h**^(*τ*)^ indicates a large weight of the corresponding criterion.

**Table 2.**
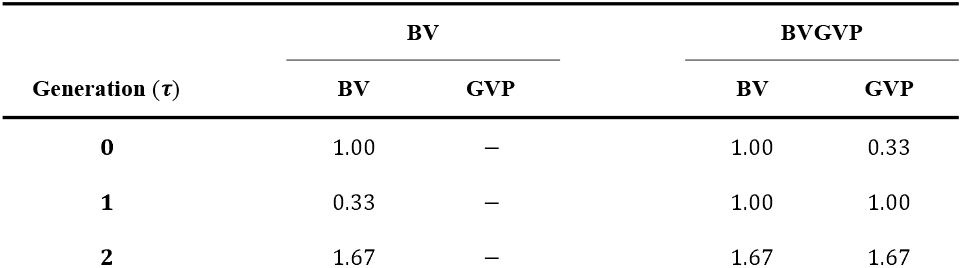
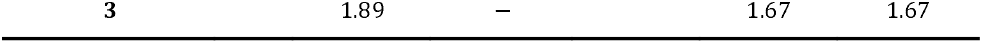
Optimized weighting parameters **h** ^(*τ*)^in different generations *τ* = 0,…, *T* − 1 for BV and BVGVP strategies.

As for the strategies without GVP, the weighting parameters in generation *τ*= 1, **h** ^(1)^showed the smallest values among all generations, and the weights in the latter generations increased through the final generation (Table 2 and Tables S1 and S2). Using strategies with GVP, the weighting parameters for the selection criteria (BV and WBV) showed higher values in generation *τ*= 1than in other generations. The tendency of the selection criteria to increase in later generations was unchanged in the strategies with GVP, and the weights for GVP were similar to those of other selection criteria across generations.

### 3.7 Optimized progeny allocation in generation *t* =0

Finally, we present the optimized number of progenies allocated to each mating pair in generation *t* =0, **b**^(0)^. Here, we compared **b**^(0)^in breeding schemes based on the equal and the six optimized allocation strategies (Table S3). The trend of which cross is more preferred was almost the same between the six optimized strategies, but the trend of how progenies are allocated to crosses was quite different. For example, the optimized strategies without GVP, such as WBV and TBV, tended to allocate more progenies to the best pair (G0_0669 x G0_0669). In contrast, the strategies with GVP, such as BVGVP and WBVGVP, tended to allocate progenies to many pairs so that the genetic diversity was maintained. From the results, we can see which mating pair was more emphasized in each strategy in not discrete but continuous ways. Thus, even if it is not practical for real breeders to carry out the exact allocation proposed by the AI breeder, the information on the optimized continuous progeny allocation will help breeders select appropriate mating pairs.

## 4 Discussion

In this study, we utilized the StoSOO algorithm to optimize the allocation of progenies and obtained quasi-optimal solutions for the set of weighting parameters **h**^(*τ*)^given a finite number of function evaluations. Utilizing these quasi-optimized weighting parameters for the allocation strategy can be justified because the true genetic gain corresponding to each set of parameters almost increased as StoSOO conducted more function evaluations (Figure S2). Thus, when applying this novel optimization approach to actual breeding schemes, the appropriate number of function evaluations will be determined according to the computational resources, and within that number of evaluations, the quasi-optimized parameters obtained from StoSOO will be used to allocate progenies.

After optimizing the weighting parameters using StoSOO, we compared the optimized allocation strategies with equal allocation strategies. The simulation results showed that the final genetic gain was higher in the following order: optimized strategies with GVP (BVGVP, WBVGVP, TBVGVP), optimized strategies without GVP (BV, WBV, TBV), and non-optimized strategy (EQ) (Figure 2A, C). These results suggest that the developed optimization framework for the progeny allocation is efficient in conventional GS and that considering the genetic diversity for later generations in the optimization framework can further improve the final genetic gains even in mid-term breeding programs (*T* = 4). In particular, the superiority of the optimized strategies with GVP was significant because those strategies guaranteed a certain level of the final genetic gains compared to other strategies, even when random factors, such as recombination between markers and segregation of alleles, are not realized in a real breeding scheme as they are in the optimization process (Figure 2B). Thus, because the optimized strategies with GVP are expected to consistently produce better genotypes than conventional GS, breeders can easily adopt our novel optimization framework in actual breeding schemes. Also, even when breeders find it difficult to implement the progeny allocation strategy proposed by the AI breeder as it is in actual breeding schemes, they can use progeny allocation results to select mating pairs as the results provide the information on which cross is likely to contribute to the production of better genotypes in the future generation (Table S3).

On the other hand, the impact on the final genetic gain caused by the type of criteria (BV, WBV, or both) used for allocation was much smaller than that caused by the existence of GVP (Figure 2A, C). Because WBV was always utilized in the selection step in this study, we succssfully maintained the genetic diversity throughout the breeding scheme regardless of the allocation strategy (Figure 2D), which may reduce the superiority of WBV in the allocation step against BV. We do not show the results here, but when BV was utilized in the selection step, the final genetic gains drastically decreased compared to when WBV was utilized for selection, regardless of the allocation strategy. Thus, since the choice of selection criteria in the selection step has a more significant impact on the final genetic gain than in the allocation step, it is strongly recommended that WBV instead of BV be used in the selection step when applying the developed method to a breeding scheme.

When focusing on the genetic gains in *t* = 1, the optimized strategies with GVP showed lower genetic gains than those without GVP (Figure 2A). In contrast, the optimized strategies with GVP showed larger genetic variances than those without GVP from *t* = 1 (Figure 2D). These results suggest that the optimized strategies with GVP were able to attain higher genetic gains in the final generation by their success in maintaining genetic diversity throughout the schemes and converting it into genetic gain toward the final generation, even in mid-term breeding programs (*T* = 4). When focusing on the optimized parameters for the strategies with GVP, the optimized weights became larger after *t* = 2, which meant that the selection intensity automatically became higher in later generations (Table 2). In addition, when focusing on the optimized weighting parameters for the strategies without GVP, the optimized weight for BV in *t* = 1was smaller than that for WBV. This may be because the larger weight of BV in the earlier generation led to less diversity, and the optimizer controlled the selection intensity by adjusting the weight to avoid such a situation. Thus, the developed framework to optimize the allocation strategy of progenies can automatically adjust the weights in each generation in an interpretable form for breeders and is expected to deal with various situations, such as breeding programs with varying deadlines *T*, by estimating proper weights for each selection criterion. In other words, although we assumed the situation close to small-scale plant breeding schemes with a relatively small number of breeding cycles and a small population size as one example of the simulation settings in this study, we can easily apply our novel framework to larger-scale plant breeding programs, and further animal breeding programs, by considering the assumed conditions such as mating constraints and changing the simulation settings proceeded by the AI breeder. Also, although we focused on the allocation strategies to prove that our future-oriented simulations can optimize the decision-making in breeding, we can further extend our framework to consider the optimization of other strategies, such as selection intensity or selection criteria used in the selection step by introducing parameters regarding those strategies to the simulation settings.

The developed future-oriented simulation-based framework achieved much higher final genetic gains than the GS method without optimized allocation of progenies. In particular, the final genetic gains for TBVGVP improved by more than 9 % on average compared with those for the non-optimized equal allocation strategy (Table 1). This improvement was quite significant for the genetic gains in just four generations because we compared the developed framework to the GS based on WBV (Goddard, 2009; Jannink, 2010), one of the most promising selection strategies in mid- and long-term breeding programs to date, not phenotypic selection or conventional GS based on BV (Meuwissen et al., 2001). Also, since the optimization algorithm was forced to terminate in the middle for the results of the ten replicates for the phenotype simulation due to the computational limitation, we may still underestimate our optimized results and can expect higher improvement than that obtained in Table 1. However, it is important to note that we used true marker effects to compute the selection criteria and GVP in this study, as the previous studies proposing the selection criteria did (Kemper et al., 2012; Daetwyler et al., 2015; Goiffon et al., 2017; Müller et al., 2018; Moeinizade et al., 2019). In actual breeding schemes, we cannot know such true marker effects and must use estimated marker effects based on GP models. Because the optimization algorithm developed in this study can be largely influenced by the estimation accuracy of GP models, developing a robust method to optimize the allocation strategy in such cases is crucial in future studies. The other important factor when assuming real breeding schemes is the timing of GP model updates. If we can appropriately update the GP model, the estimation accuracy of the marker effects and optimization performance of the allocation strategies will be improved, which will directly lead to further improvement in the final genetic gains. In addition, we can deal with long breeding schemes by conducting appropriate model updates. Thus, we will further investigate the potential of our optimization framework by assuming real breeding schemes with the factors discussed above.

Genomic selection is a promising technique that contributes to the acceleration of breeding. In this study, we introduced and developed a novel framework that can upgrade conventional GS in breeding schemes by optimizing the allocation strategy of progenies via future-oriented simulations. To optimize the allocation strategy, we parameterized the strategy via softmax conversion by utilizing the selection criteria and expected genetic variance of progenies. From the simulation results, our novel framework with the optimized allocation significantly outperformed the non-optimized strategies, especially when we added the genetic variance of progenies to the algorithm. Thus, this future-oriented optimization framework for breeding schemes can contribute to the acceleration, high efficiency, and optimization of plant breeding at the hybridization stage, which will lead to the optimization of various kinds of decision-making in plant and animal breeding.

## DATA AVAILABILITY STATEMENT

All data were simulated in this study. Because we performed many breeding simulations, we cannot share all datasets themselves. Instead, we share our scripts, including the simulation of data. The scripts used in this study are available from the “KosukeHamazaki/SCOBS” repository on GitHub, https://github.com/KosukeHamazaki/SCOBS. In this study, we utilized the R package “myBreedSimulatR” to simulate and optimize breeding schemes. “myBreedSimulatR” is available from the “KosukeHamazaki/myBreedSimulatR” repository on GitHub, https://github.com/KosukeHamazaki/myBreedSimulatR. This package is a variant of “breedSimulatR”, which is available from the “ut-biomet/breedSimulatR” repository on GitHub, https://github.com/ut-biomet/breedSimulatR.

## AUTHOR CONTRIBUTIONS

KH implemented the simulation of breeding schemes, developed the method for optimizing allocation strategies in breeding schemes, and drafted the manuscript. KH and HI conceived and designed the study. HI provided administrative support and supervised the study. All authors have read and approved the final manuscript.

## FUNDING

This study was supported by the Japan Society for the Promotion of Science (JSPS) KAKENHI Grant Number JP 22H02306. The funders had no role in the study design, data collection, analysis, decision to publish, or preparation of the manuscript.

## ACKNOWLEDGMENTS

We are grateful to Mr. Julien Diot, a developer of the “breedSimulatR” package, for fruitful discussions on the implementation of breeding simulations. We would like to thank Dr. Yohei Akimoto for fruitful discussions on how to optimize decision-making in a breeding scheme. We are also grateful to Dr. Bo Zhang for her valuable comments when revising our manuscript. We would like to thank the Japan Society for the Promotion of Science (JSPS) for funding our study. We would also like to thank Editage (www.editage.com) for English language editing.

## CONFLICT OF INTEREST STATEMENT

Not applicable. The authors declare that they have no competing interests.

## SUPPLEMENTAL DATA

The Supplementary Material for this article can be found online at:

## Notes

https://github.com/KosukeHamazaki/SCOBS

https://github.com/KosukeHamazaki/myBreedSimulatR

### Competing Interest Statement

The authors have declared no competing interest.

### Summary of Updates

All the manuscripts were revised regarding the concept of this study. Especially, we added a clearer explanation of the concept and background of the study and why we adopted the simulation settings.

