## Supplementary Information for "AI-assisted selection of mating pairs through simulation-based optimized progeny allocation strategies in plant breeding"

![frontiers]

### 1 Supplementary Figures and Tables

#### 1.1 Supplementary Figures

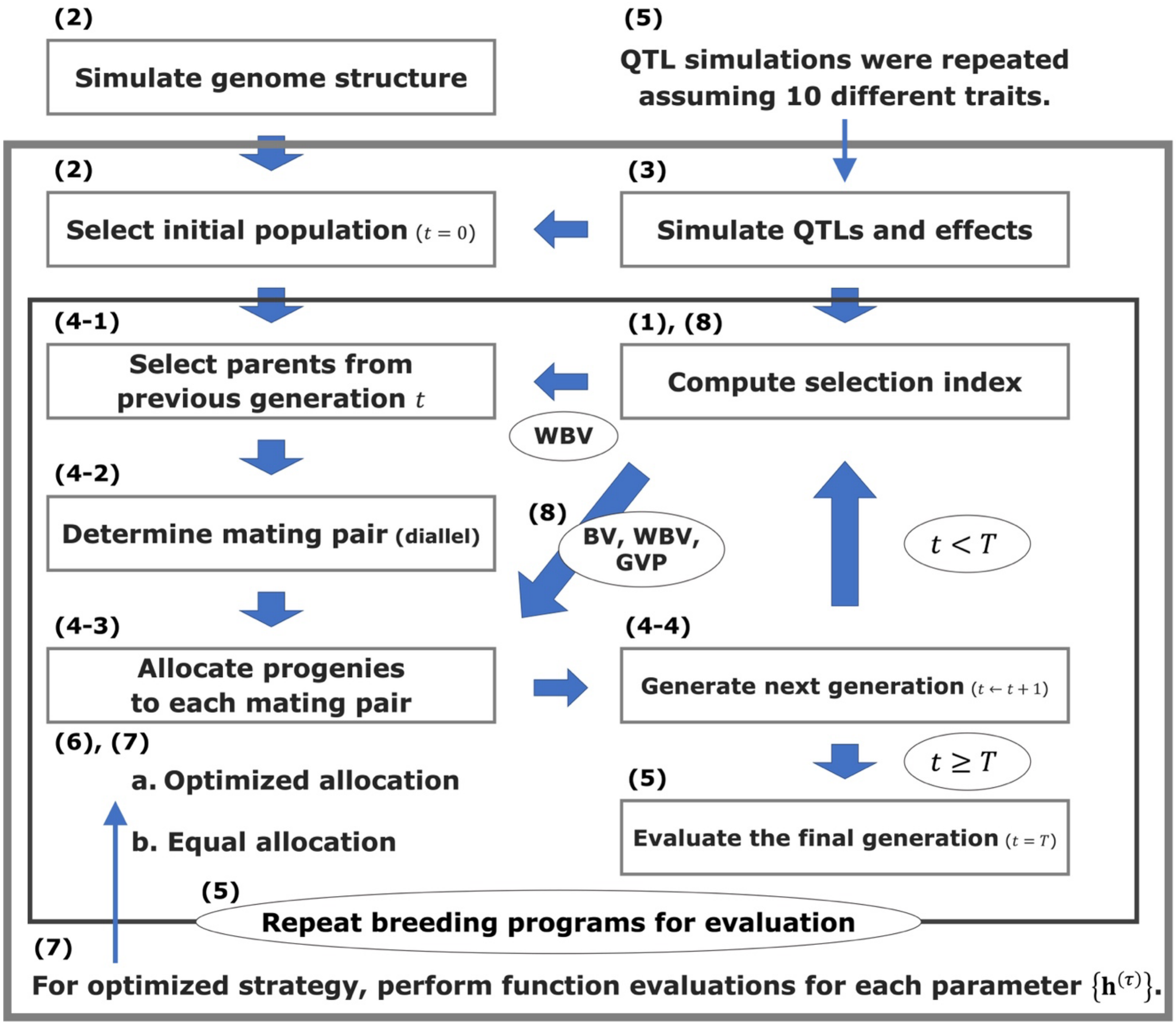


**Supplementary Figure 1.** Flowcharts for the overview of our simulation study. The numbers in parentheses indicate the order in which they will be explained.

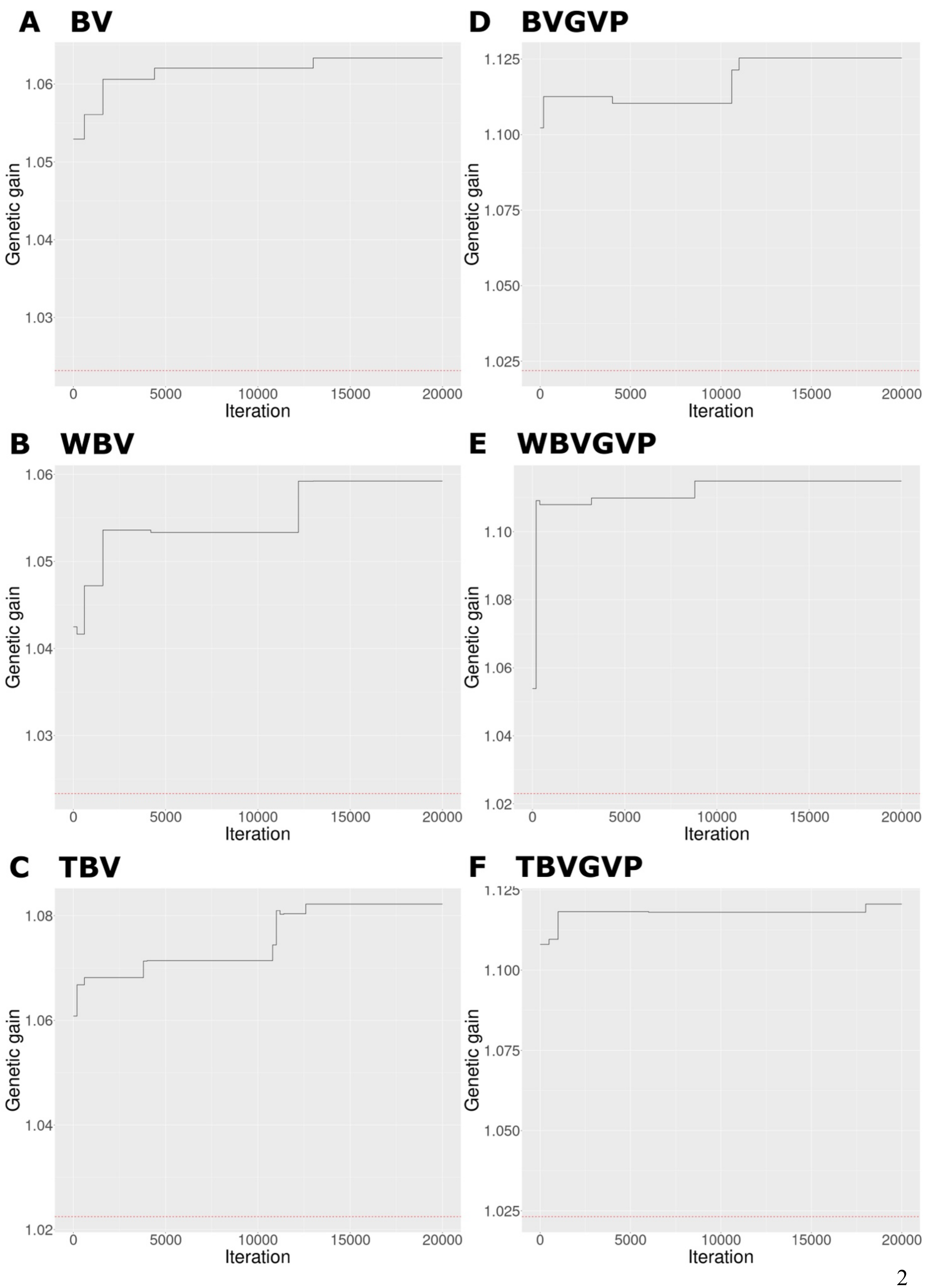
A BV
B WBV
C TBV
D BVGVP
E WBVGVP
F TBVGVP
Genetic gain
Iteration
0
5000
10000
15000
20000
1.02
1.03
1.04
1.05
1.06
1.08
1.10
1.025
1.050
1.075
1.100
1.125

**Supplementary Figure 2.** Change in function values for each allocation strategy optimized by the StoSOO algorithm. The black solid line shows the change in function values (expected final genetic gains evaluated by 10,000 simulations) for each allocation strategy optimized by the StoSOO algorithm. The red dashed line shows the expected final genetic gain based on the equal allocation strategy. Each subplot corresponds to the result for each optimized strategy as follows; (**A**) BV, (**B**) WBV, (**C**) TBV, (**D**) BVGVP, (**E**) WBVGVP, (**F**) TBVGVP. The abbreviation of each strategy is as same as that in Figure 2.

### 1.2 Supplementary Tables

**Supplementary Table 1.** Optimized weighting parameters $\mathbf{h}^{(\tau)}$ in different generations $\tau = 0, \dots, T-1$ for WBV and WBVGVP strategies.

| | **WBV** | | **WBVGVP** | |
|---|---|---|---|---|
| **Generation ($\tau$)** | WBV | GVP | WBV | GVP |
| **0** | 1.22 | – | 0.33 | 0.33 |
| **1** | 0.85 | – | 1.00 | 1.00 |
| **2** | 1.88 | – | 1.67 | 1.67 |
| **3** | 1.67 | – | 1.00 | 1.00 |

**Supplementary Table 2.** Optimized weighting parameters $\mathbf{h}^{(\tau)}$ in different generations $\tau = 0, \dots, T-1$ for WBV and TBVGVP strategies.

| | **TBV** | | | **TBVGVP** | | |
|---|---|---|---|---|---|---|
| **Generation ($\tau$)** | BV | WBV | GVP | BV | WBV | GVP |
| **0** | 1.00 | 0.33 | – | 1.00 | 0.33 | 0.33 |
| **1** | 0.33 | 1.00 | – | 1.00 | 0.33 | 1.00 |
| **2** | 1.89 | 1.00 | – | 1.67 | 1.00 | 1.00 |
| **3** | 1.67 | 1.67 | – | 1.00 | 1.00 | 1.00 |
